# Variation in Head and Pinna Morphology of Preserved *Peromyscus Spp*. Specimens and Implications for Auditory Function

**DOI:** 10.1101/2024.12.12.628269

**Authors:** Casey E. Sergott, Katelynn Rodman, Nathaniel T. Greene, Ben-Zheng Li, Genesis A. Alarcon, Fabio A. Machado, Elizabeth A. McCullagh

## Abstract

The characteristics of an animal’s head and pinna mark the beginning of auditory communication. Auditory communication is broadly achieved by receiving sounds from the environment and plays a vital role in an animal’s ability to perceive and localize sounds. Natural history museums and collections along with their vast repositories of specimens provide a unique resource for examining how the variability in both the size and shape of the head and pinna cause variability in the detection of acoustic signals across species. Using this approach, we measured the dimensions of the head and pinna of over 1,200 preserved specimens of *Peromyscus boylii*, *P. californicus*, *P. gossypinus*, *P. leucopus*, *P. maniculatus*, and *P. truei*, followed by a series of head-related transfer functions (HRTFs) on several individuals to study the relationship between morphology and available auditory information. Our morphological results show significant variation in pinna length and width, as well as in the distance between the two ears across the six species. ITDs and ILDs were calculated and demonstrated consistent results across species, suggesting the differences in head and pinna size do not significantly modify these cues. Not only does this study contribute to existing research on external morphology and auditory function, but it also provides valuable insight into the use of preserved specimens in auditory research, an area that is currently understudied.

**Summary statement:** This work aims to provide insight into using natural history museum specimens for morphological research pertaining to the auditory system in small mammals.

## Introduction

Morphological features give great insight into the function of the structures that comprise an animal. For example, the skeleton, dentition, and body size/composition tell us much about an animal’s locomotion, habitat, diet, and even behavior (Schwab et al., 2011). Similarly, the head and pinna are the first contact points for sound waves before reaching the inner ear. The pinna functions primarily to localize and direct sounds into the middle and inner ear, where they can then be interpreted as electrical signals in the brain. Though it is also thought that pinnae play a role in thermoregulation in many species, an idea first proposed by biologist Joseph Asaph Allen in 1877, which predicts that animals living at higher latitudes (generally colder environments), should have smaller appendages (i.e., limbs, pinna, or tail) than animals living in lower latitudes based on the properties of heat conservation (Allen, 1877; Fan et al., 2019; Alhajeri et al., 2020). Tracking these different pressures on morphology requires ample samples covering a broad geographic and temporal range, such as those present in zoological collections.

Natural history museums and zoological collections are permanent repositories of biodiversity throughout history and provide immense value to scientific research. Collections often work directly in collaboration with zoos and research institutions to allow their specimens to be used for studying evolution, morphology, conservation, genetics, and much more across a variety of scientific disciplines (Nakahama, 2020; Poo et al., 2022). They are often overlooked in auditory research as many of the structures of the auditory system are small, fragile, located within the skull, and are therefore not easily preserved (Stoessel et al., 2016). Limited research has been conducted, however, on the overall pinna size and shape (such as here), the tympanic membrane, ossicle sizes, and middle ear cavities (Stephens et al., 2015, Mason et al., 2017). Thus, to expand our understanding of the usefulness of museum collections in auditory research, we need to study specimens from a group of species that are variable in both their morphology and habitats, such as deer mice from the genus *Peromyscus* (Cricetidae; Rodentia).

Deer mice are among North America’s most abundant groups of animals (Bedford and Hoekstra, 2015). Many species, such as *P. leucopus* and *P. maniculatus,* are colloquially regarded as habitat generalists, occupying diverse habitats, including grasslands, mountains, and forests. In contrast, some species, such as *P. truei* and *P. californicus* occupy more specialized regions, such as pinyon-juniper woodlands and damp oak-woodlands, respectively (Fellers, 1994; Kobrina et al., 2021). Here, we focused on six *Peromyscus* species – *P. leucopus*, *P. maniculatus*, *P. boylii*, *P. truei*, *P. gossypinus*, and *P. californicus* to assess morphological and functional differences in head and pinna size. These species were selected due to the habitats in which they occur, giving insight into whether geographic location plays a role in terms of either form or function. *P. leucopus* and *P. maniculatus* were chosen because they are known for being some of the most widespread *Peromyscus* species, and there are only a few areas of North America that they do not inhabit. *P. truei*, on the other hand, are primarily found on the west coast of North America and are known for thriving within pinyon-juniper woodlands but are also known to occupy rocky slopes and grasslands. *P. boylii* thrive at higher elevations (above 2,000 m), especially throughout mountainous regions in the Western U.S. (Ribble et al., 2002). *P. gossypinus* and *P. californicus* were selected based on their intermediate environmental preferences, providing a comprehensive representation across habitats. If there are auditory morphology differences within *Peromyscus*, we expect to find them within this group of species.

The biomechanics of hearing and sound localization encompasses the highly complex interplay between the ear and brain structures. The ability to accurately locate and process complex acoustic stimuli from the environment has been linked to the pinna in multiple mammalian species, including bats (Aytekin et al., 2004), guinea pigs (Greene et al., 2014), cats (Young et al., 1996; Xu and Middlebrooks, 2000; Rebillat et al., 2014; Benichoux et al., 2016), and chinchillas (Heffner et al., 1996, Osmanski and Wang, 2001; Koka et al., 2011). Particularly, the presence of the pinna plays a major role in the front/back and vertical discrimination of acoustic signals (Heffner, Koay, and Heffner, 1996; Jones et al., 2011; Alves-Pinto et al., 2014). This is due to the folds and convolutions of the pinna which filter sounds to create unique spectral patterns (notches and peaks) that the brain then associates with specific locations in elevation (Rice et al., 1992; Middlebrooks, 2015; Anbuhl et al., 2017). After the pinna receives a sound, the brain can then assess the approximate location in azimuth, relying on the computation of two cues: interaural time differences (ITDs) and interaural level differences (ILDs) or the shift in time and sound level of a particular cue at each of the pinnae (Wightman and Kistler, 1993; Hartmann, 2021). To resolve front/back ambiguity, the position and orientation of the head, body, and pinna, must be used due to the similarity in ITD and ILD of sounds arriving from the front and back. Head-related transfer functions (HRTFs) are transfer functions that account for the size and shape of these external features, to determine how a specific sound stimulus is both received and impacted by the pinna of an individual at a set distance and elevation in space in the ear canal (Lei and Peissig, 2020). They are unique to every individual and lend perspective into whether morphological differences are correlated with functional differences in signal detection (Pec et al., 2007). As the acoustics of the HRTF rely only on anatomical/morphological features of the head and ear, they can easily be performed on preserved zoological specimens (Rebillat et al., 2014; Benichoux et al., 2016).

Importantly, this research demonstrates the utility of zoological specimens in studying the impacts of head and pinna size differences on auditory cues available to different species. Furthermore, our work helps to uncover the relationship between pinna morphology, habitat, and auditory function using preserved specimens of the six *Peromyscus* species mentioned. We expect that if differences are seen in head and pinna morphology, then sound localization cues may also be affected, reflecting different auditory niches. We also aim to test the accuracy of using zoological specimens in morphological research by performing statistical analyses on new and old measurements, in which we expect to find shrinkage in the new measurements due to the drying time and age of the specimens.

## Materials and methods

### Animal models

Preserved specimens of *Peromyscus boylii*, *P. californicus*, *P. gossypinus*, *P. leucopus*, *P. maniculatus*, and *P. truei* were provided by the Oklahoma State University’s Collection of Vertebrates. Individual specimens were given to the collection through various institutional researchers and zoological donations over the years, with some specimens used dating back to 1903. Each animal used in this study had the following data associated with them: genus, species, date of collection, precise location of collection site, name of the collector and specimen preparator, as well as initial measurements of the body, tail, hindfoot, and ear lengths. Given the variability in specimen preparation, only the best-preserved specimens were chosen for analysis (i.e., pinnae tissue was not folded or visibly damaged, and the head was stuffed to mimic the natural position as best as possible). All specimens used were collected from North America, with the majority being from Oklahoma and Colorado.

### Morphology

We measured the head and pinna of 1,274 preserved *Peromyscus* specimens (*P. leucopus*, n = 780; *P. maniculatus*, n = 382; *P. boylii*, n = 63; *P. truei*, n = 33; *P. gossypinus*, n = 12; *P. californicus*, n=4). Measurements included pinna size (length and width), distance between the pinnae (abbreviated as IT), distance from the tip of the nose to the midpoint between the pinnae (abbreviated as NT), and the effective diameter (ED) (Fig. 1). The effective diameter was calculated using the formula 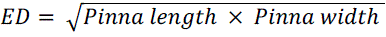 to draw inferences on sound detection capabilities. Length measurements of the pinna were taken as the maximum distance from the base to the tip, while width measurements were taken at the widest part of the pinna to obtain the greatest perpendicular distance across the structure. IT measurements were taken as the distance between ear canals, while NT measurements were taken as the linear distance from the tip of the nose to the midpoint between the two ears, measured with calipers positioned above the head. Although these measurements lacked distinct anatomical landmarks as anchor points, all measurements were performed at the same relative positions on each specimen by two researchers using the same 6-inch electronic vernier caliper (DIGI-Science Accumatic digital caliper, Gyros Precision Tools, Monsey, NY, USA) each measurement session. Due to the age and nature of zoological specimens, we also tested the accuracy of research collection measurements by comparing the new measurements to the ones taken at the time of collection. This limitation was addressed by testing the correlation between original length measurements (the only pinna measurement taken at the time of collection) and new length measurements (taken presently). All pinna measurements were taken on the right side of the animal. Mean values for each measurement type were calculated in RStudio (version 4.2.1) (Tables 1, 2).

**Figure 1.**
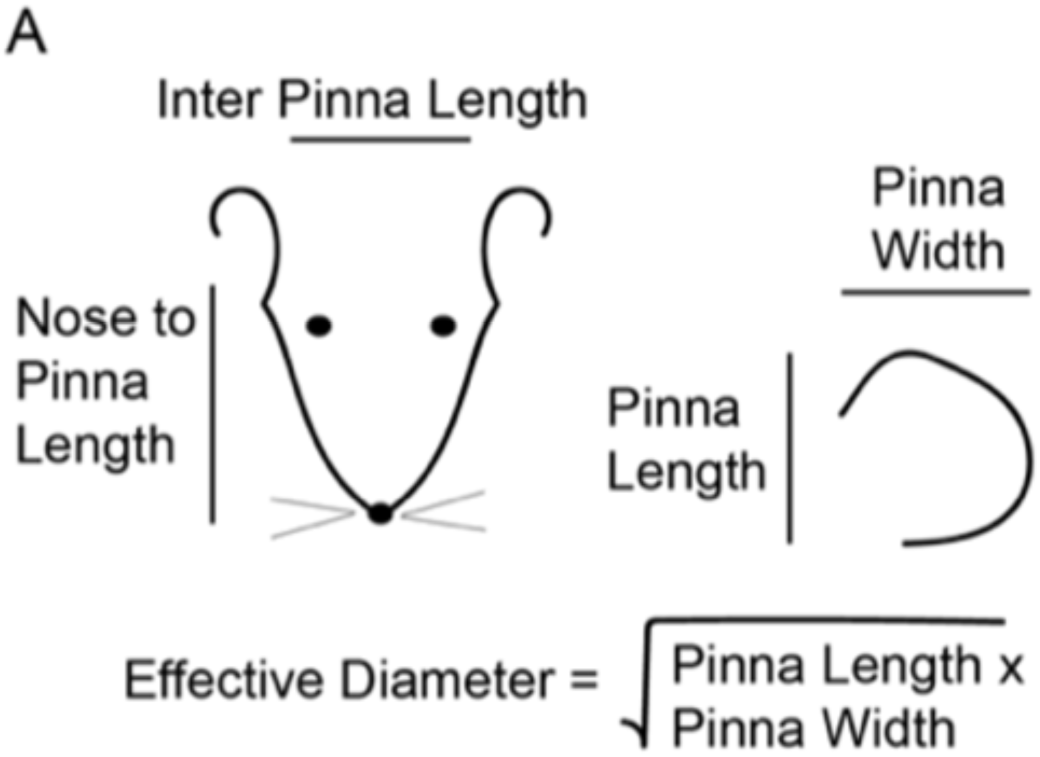
Portrayal of morphological measurements taken. A simplified depiction of the morphological measurements taken on each animal subject.

**Table 1.** Mean measurements for pinna length and width by species.

| Species | *n* | Mean length (mm) | s.d. | Mean width (mm) | s.d. |
| --- | --- | --- | --- | --- | --- |
| leucopus | 780 | 14.01 | 1.51 | 8.43 | 1.17 |
| maniculatus | 382 | 13.03 | 2.14 | 8.11 | 1.24 |
| gossypinus | 12 | 15.70 | 1.71 | 9.92 | 1.13 |
| truei | 33 | 21.67 | 2.38 | 12.49 | 1.92 |
| boylii | 63 | 17.36 | 1.26 | 10.32 | 1.08 |
| californicus | 4 | 20.14 | 1.18 | 13.40 | 2.53 |

**Table 2.**
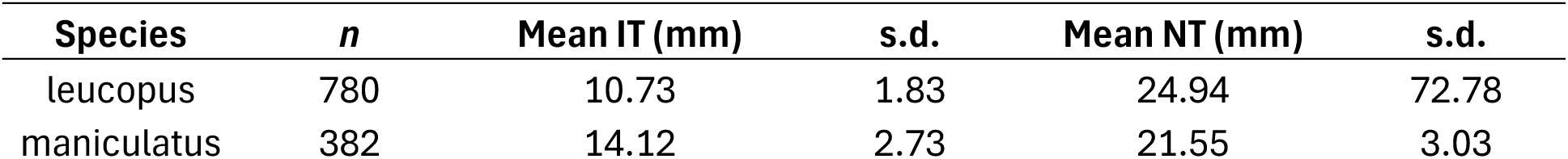
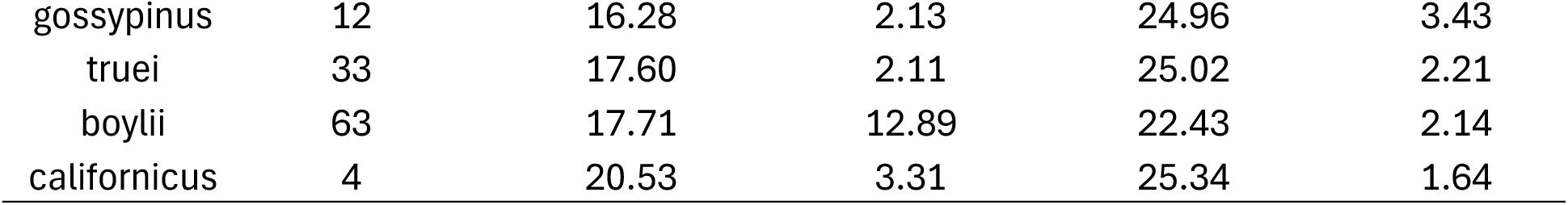
Mean measurements for inter-pinna distance and nose-to-pinna distance by species.

| Species | *n* | Mean IT (mm) | s.d. | Mean NT (mm) | s.d. |
| --- | --- | --- | --- | --- | --- |
| leucopus | 780 | 10.73 | 1.83 | 24.94 | 72.78 |
| maniculatus | 382 | 14.12 | 2.73 | 21.55 | 3.03 |
| gossypinus | 12 | 16.28 | 2.13 | 24.96 | 3.43 |
| truei | 33 | 17.60 | 2.11 | 25.02 | 2.21 |
| boylli | 63 | 17.71 | 12.89 | 22.43 | 2.14 |
| californicus | 4 | 20.53 | 3.31 | 25.34 | 1.64 |

### Head-related transfer functions

To assess auditory function in preserved specimens of *Peromyscus,* we collected a series of HRTFs to determine how sounds are captured at a specific location in the ear canal and whether morphological differences across species lead to functional changes. We conducted a series of HRTFs on a total of 18 preserved *Peromyscus* specimens (*P. leucopus* = 5, *P. maniculatus* = 5, *P. boylii* = 5, *P. truei* = 3). These specimens displayed measurements characteristic of the range observed in the means recorded from the museum sample. HRTFs were conducted in a sound-attenuating chamber (86 x 112 x86 in.; O’Neill Engineered Systems, Hartland, WI) with a custom-made speaker system array that consisted of a single movable speaker (Dayton Audio ND65-4, Springboro, OH), capable of moving 180° around the subject in 10° increments, making for a total of 19 possible locations on the horizontal plane and 19 possible locations on the vertical plane for a total of 361 potential measurements (Fig. 2). In azimuth, –90° and +90° were defined as directly on the left and right sides, respectively, and 0° being directly in the middle. In elevation, 0° V was defined as directly in front of the subject, with 180° V being directly behind the subject.

**Figure 2.**
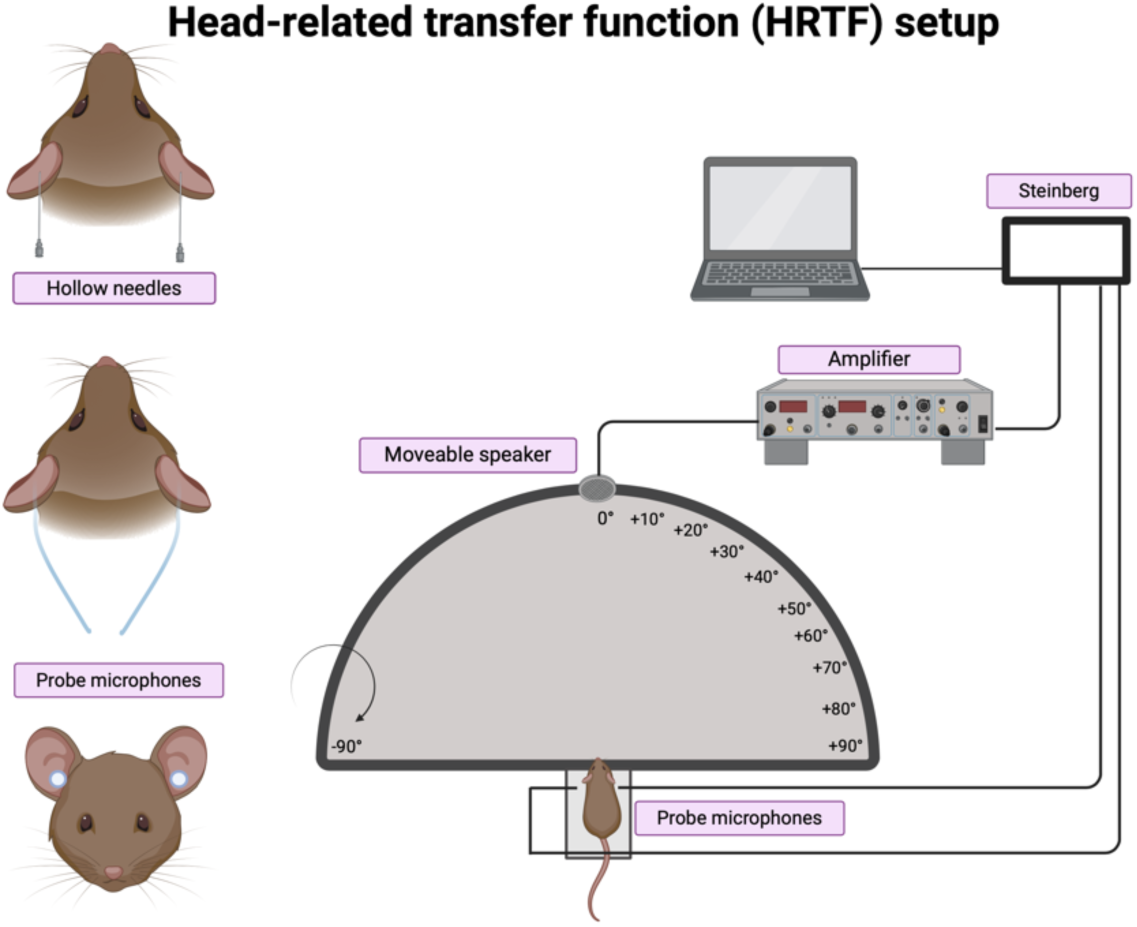
Head-related transfer function equipment. Figure representation of the custom-made HRTF setup and microphone placement strategy used in this study. The array offers a total of 361 unique possible speaker locations along the horizontal and vertical axes.

In light of the logistical and technical constraints of the experimental setup, we focused only on capturing ITD and ILD data. We, therefore, took measurements at each location on the horizontal axis (–90° to +90°) both in front of and behind the animal, for a total of 38 measurements. ER-7C probe tube microphones (Etymotic Research Inc., Elk Grove Village, IL) were placed into the preserved specimen’s ear canal by threading the tubing through a hollow 14-gauge needle carefully inserted through the base of the pinna, remaining as close to the eardrum as possible, ensuring the microphone tips did not come in contact with any tissue or hair. Due to the fragility of zoological specimens, small perforations were made with the tip of the needle near the insertion point to eliminate tearing and excess space around the microphones during the placement (Fig. 2). Measurements were taken with room acoustics software REW (Room EQ Wizard, version 5.20.13 Pro Upgrade, Montrose, Angus, Scotland, UK). A Steinberg UR22C (Steinberg Media Technologies GmBH, Hamburg, Germany) was used as the audio interface with a Sony stereo amplifier (STR-DH190, New York, NY) to drive the speaker. The system was calibrated using microphone calibration files provided by Etymotic as well as by performing a manual calibration using a dB meter and sound card to ensure the input and output levels were within 6dBfs (full scale) of each other. To eliminate any loudspeaker effects, additional calibration measurements were taken with only the microphones positioned on the platform in a similar place to where the microphones would rest with the animal present.

Once the initial calibration was complete, the specimen was placed on an acrylic platform in the center of the array, with the microphones precisely positioned in the ear canal. Acoustic stimuli were presented as 128k point sine sweeps (–28 dBfs), starting at 250 Hz and ending at 20 kHz over 3 seconds with a sampling rate of 44.1 kHz, using a loopback timing reference. Each sine sweep stimulus was repeated five times at each location on the horizontal axis while the microphones recorded responses from the left and right sides simultaneously. The responses were amplified and digitized with the Steinberg UR22C and displayed by the REW software as transfer function and impulse response from the FFT (Fast Fourier Transform) of the left, right, and combined signals. HRTFs were calculated as the gain of sound pressure level (SPL) between the response received with the specimen present and the microphone-only response. ITDs were derived by finding the time shift between the signals received by the left and right ear microphones that maximize the signal cross-correlation. ILDs were calculated as the frequency-dependent difference in gain between the right ear microphone relative to the left at each speaker position (in dB).

### Statistical analysis

All figures and statistical analysis were completed in RStudio (version 4.2.1) or Python (Version 3.11.7). Because preservation is thought to alter morphology by the drying out of tissues via shrinkage, a thorough analysis was taken to ensure drying time and preparation variation were accounted for in overall morphological measurements by comparing original measurements (taken at the time of collection) to the new measurements (taken presently). The correlation coefficients (R-values), as well as a 1:1 correspondence shown with a solid black line which represents what a perfect correlation between measurements would look like, are reported. To compare old and new measurements, as well as measurements within and between species, a linear mixed-effects model (LMM) and one-way ANOVA (R packages lmerTest and lme4) were used to account for random variability between specimen measurements (measurement type (length, width, etc.) was set as the response variable, species as the fixed effect, and animal number as the random effect variable). If significant differences in fixed effects were found through the model, Tukey’s Honest Significant Difference (THSD) test was then performed to show all pairwise comparisons between species. Measurements within and between species are depicted by box and whisker plots. For HRTF analysis, amplitude spectra were calculated using a 512-point fast Fourier transform (FFT) and filtered with a 1/48 octave filter within REW. HRTF traces were adjusted by a factor of 2.5 and were exported as .txt files and analyzed in Python.

## Results

### Morphology

The correlation values between the new measurements to the ones taken at the time of collection are moderate (*P. leucopus*, R = 0.26; *P. maniculatus*, R = 0.33; *P. boylii*, R = 0.41; *P. truei*, R = 0.42) (Fig. 3), but show no systematic bias for shrinkage. Due to the amount of error in the measurements, we focused on differences across species, not individuals. The LMM identified statistical differences in three measurements: pinna length, pinna width, and inter-pinna distance (IT; Fig. 4, Tables S1, 2, 3, 4). IT showed the smallest difference across species means, with a mean square between groups (MSB) of 1272.5 (F = 100.5, p = <2e-16), while pinna length had the greatest differences, with an MSB of 632.9 (F= 210, p = < 2e-16). Pinna width had an MSB of 168.66 (F = 113.6, p = < 2e-16), showing less pronounced contrasts across species. No significant difference was found between species in NT measurements, MSB = 604, F = 0.183, p = 0.969. Pairwise comparisons for each measurement across species can be found in Supplementary Tables 1, 2, 3, and 4. Overall, *P. truei* had the longest and widest pinnae and the greatest IT compared to the other species analyzed (Fig. 3). Notably, *P. leucopus* and *P. maniculatus*, two very closely related species, overlapped in measurements yet were still significantly different in pinna width and IT. *P. leucopus* had smaller ITs (p < 0.001) with larger pinna widths compared to *P. maniculatus* (p < 0.001).

**Figure 3.**
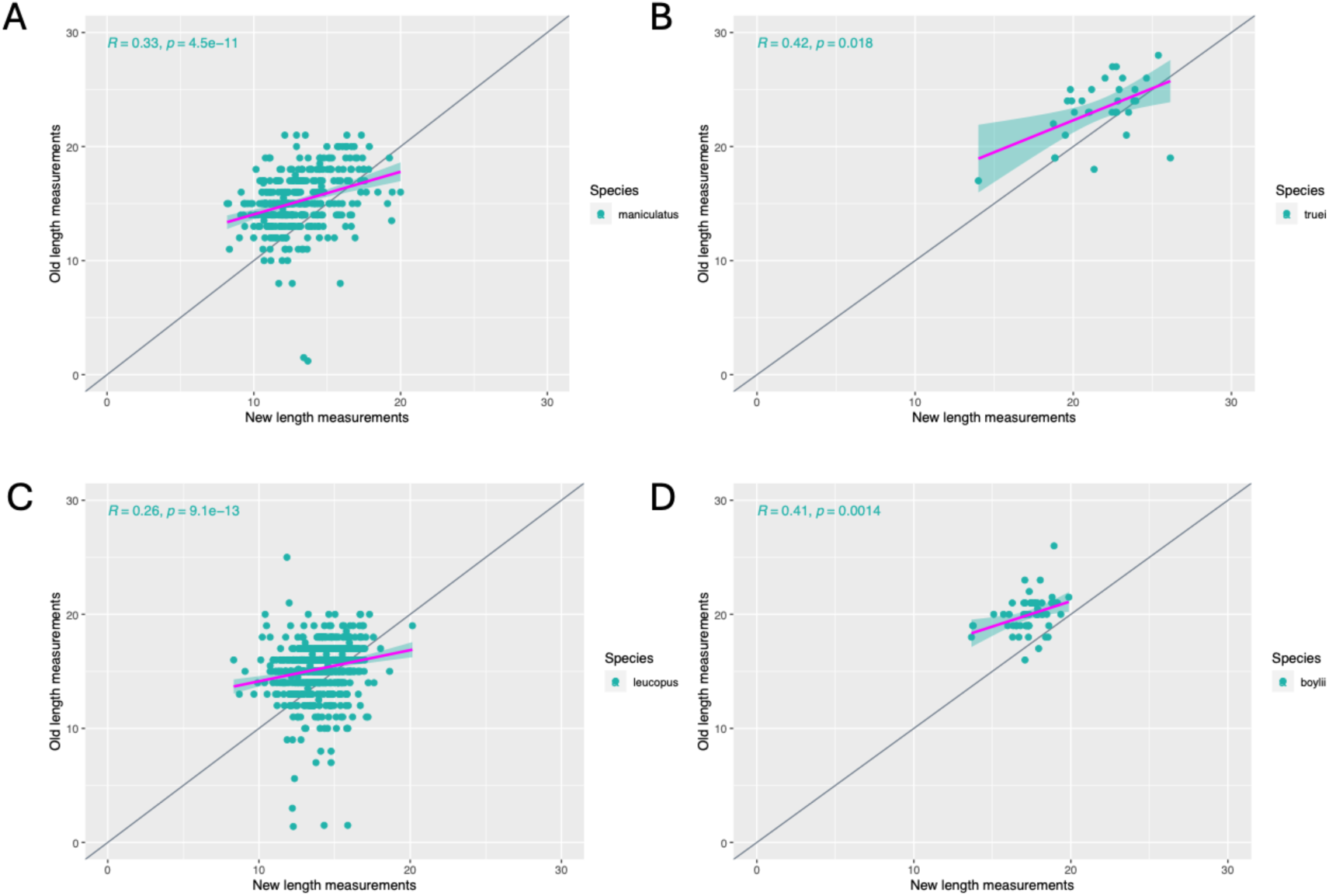
Comparison of new and old pinna length measurements. Comparison of original measurements taken at the time of collection to present measurements in *P. maniculatus* (A), *P. truei* (B), *P. leucopus* (C), and *P. boylii* (D). Each dot represents a measurement point, while the solid black line shows a 1:1 correspondence.

**Figure 4.**
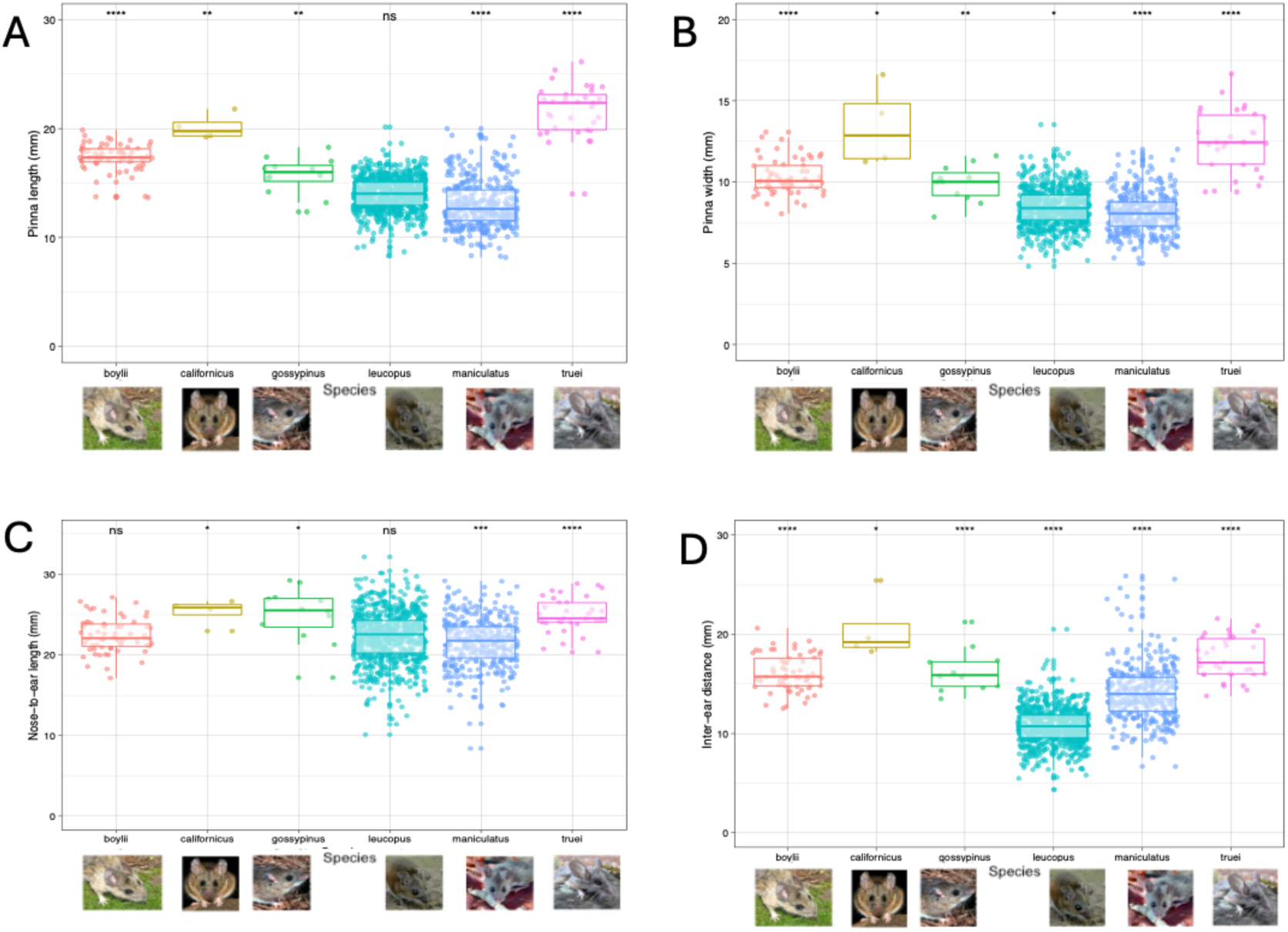
Head and pinna measurements. Pinna length (A), pinna width (B), NT (C), and IT (D) measurements in preserved specimens of *P. boylii*, *P. californicus*, *P. gossypinus*, *P. leucopus*, *P. maniculatus*, and *P. truei*. Each point represents an individual measurement. (Significance codes: p = 0 ‘***’, p = 0.001, ‘**’, p = 0.01, ‘*’, p = 0.05 ‘.’, ns = not significant)

### Head-related transfer functions

HRTFs were used to explore how morphological features of the head and pinna influence how sound cues are captured in preserved *Peromyscus* specimens. Although six species were initially considered for this study, only four had sufficient specimens available in the research collection utilized for this study. Therefore, the lack of representation led to the exclusion of P*. gossypinus* and *P. californicus* from the HRTF analysis. HRTF gain (characterized by peaks in y-axis values) and spectral notches (characterized by dips in y-axis values) specifically give insight into how different frequencies are amplified (gain) or attenuated (spectral notches) based on the morphological features of the individual.

ITD was similar for all species tested across measured angles (Fig. 5). This measurement along with ILD provides spatial acoustic information to the animal. ITD values showed consistent patterns of cue availability across all four species, with notably greater variability for locations behind the animal compared to the front. We see the largest difference in maximum ITD between *P. boylii* and *P. truei*, which does not correlate with the differences in head and pinna sizes. *P. leucopus*, despite having the smallest pinna width across species, has the second longest ITD (380.95 µs). The average maximum and minimum ITD values for each species can be found in Supplemental Table 5.

**Figure 5.**
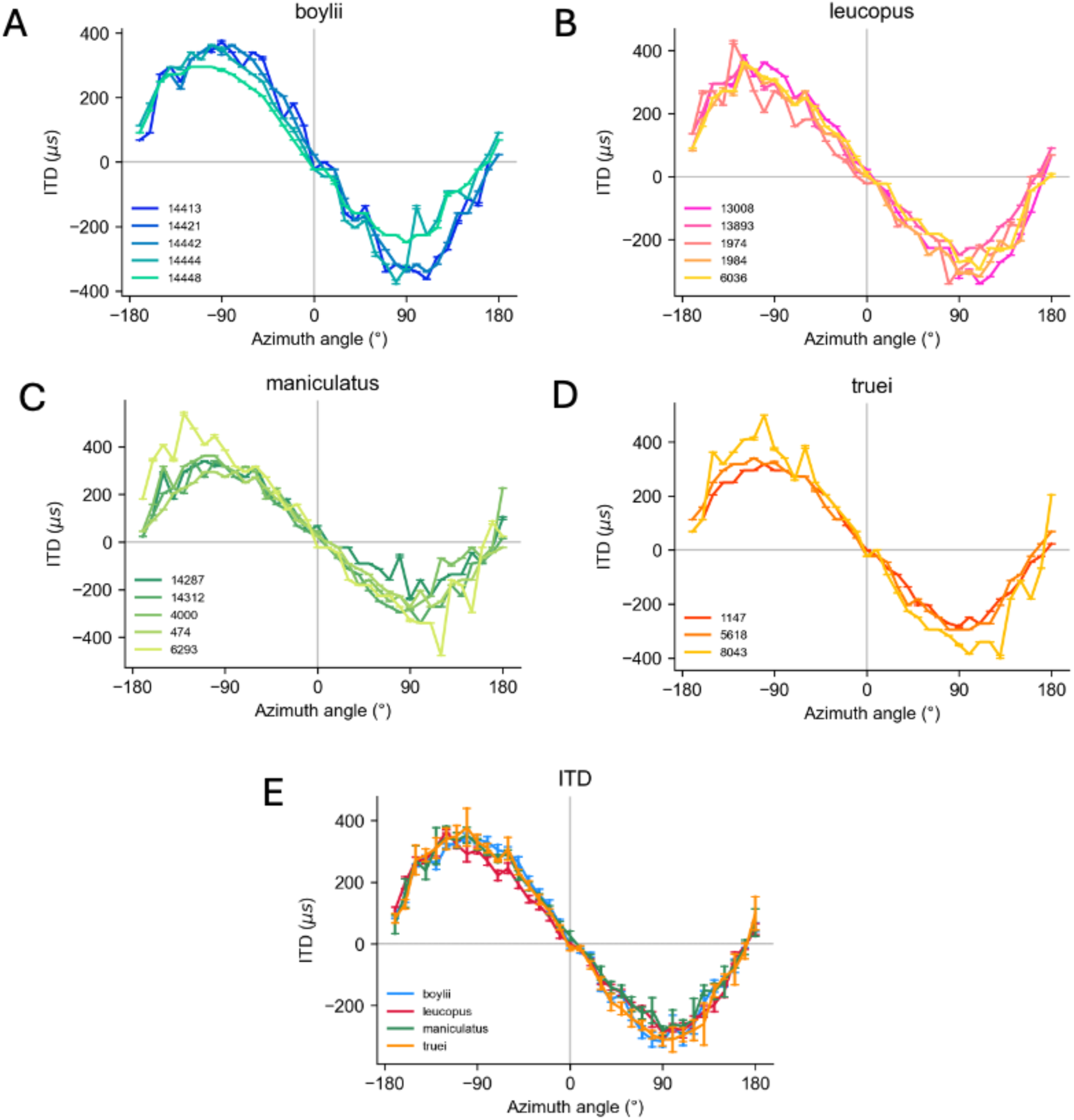
ITD results for *P. boylii* (A), *P. leucopus* (B), *P. maniculatus* (C), and *P. truei* (D). Measurements were taken in front of and behind the animal in 10° increments in azimuth. –90° to +90° azimuth represents measurements taken in front of the animal while +/-90° to +/-180° indicates measurements taken behind the animal. Each line represents a separate individual in the species in (A-D). The combined ITD results for each species are shown in (E).

As mentioned above, ILD, calculated as the difference in sound intensity at the two pinnae, is a common cue for sound localization. Our results show that ILD is consistent across all four species and is optimized at higher frequencies, such as at 8 kHz and above, where the greatest differences are seen and, therefore, the most information is derived (Figs 6, 7). At 8 kHz, *P. maniculatus* had the highest maximum ILD of 9.94 dB, though at 16 kHz, *P. boylii* had the highest maximum ILD of 8.75 dB. Variability is consistently found around 10 kHz on the same side of the head as the measurement (Fig. 6). The maximum ILD in front of the animal at both 8 kHz and 16 kHz was greater than or equal to +/-60° azimuth (Fig. 7A, C). However, this range narrows when examining stimuli directly behind the animal. The greatest differences were seen in ILDs closer to –/+30° azimuth, and the widest variability in ILD between species occurred during stimuli presented behind the animal at 16 kHz (Fig. 7). The presence of a signal around 15 kHz across animals is likely an artifact of the measurement process and is not considered to represent meaningful or biologically relevant data.

**Figure 6.**
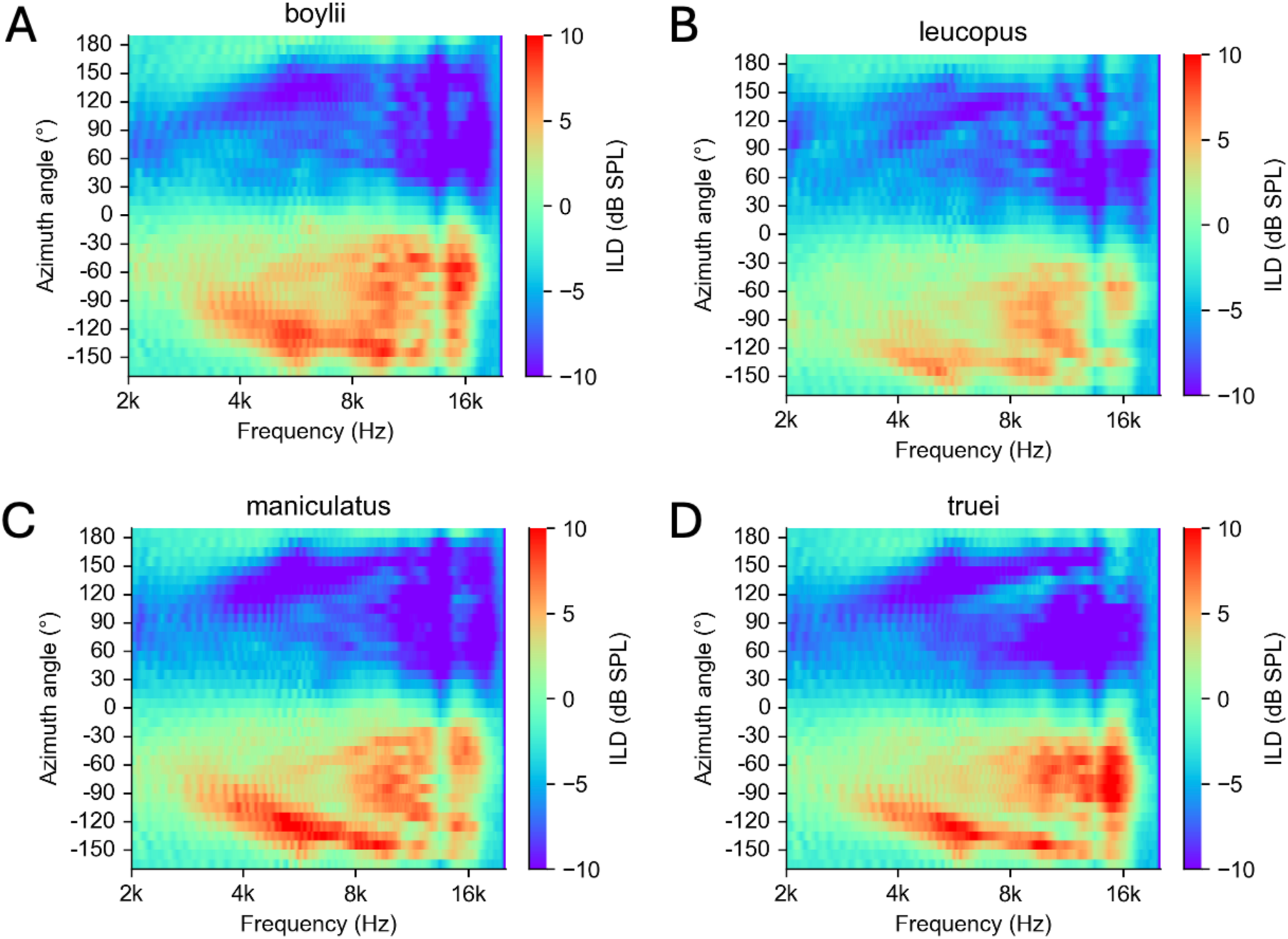
ILD spectrogram results. ILD results for P. boylii (A), P. leucopus (B), P. maniculatus (C), and P. truei (D). Measurements were taken directly in front and behind the animal at 10° increments in azimuth. –90° to +90° azimuth represents measurements taken in front of the animal while +/-90° to +/-180° azimuth represents measurements taken behind the animal.

**Figure 7.**
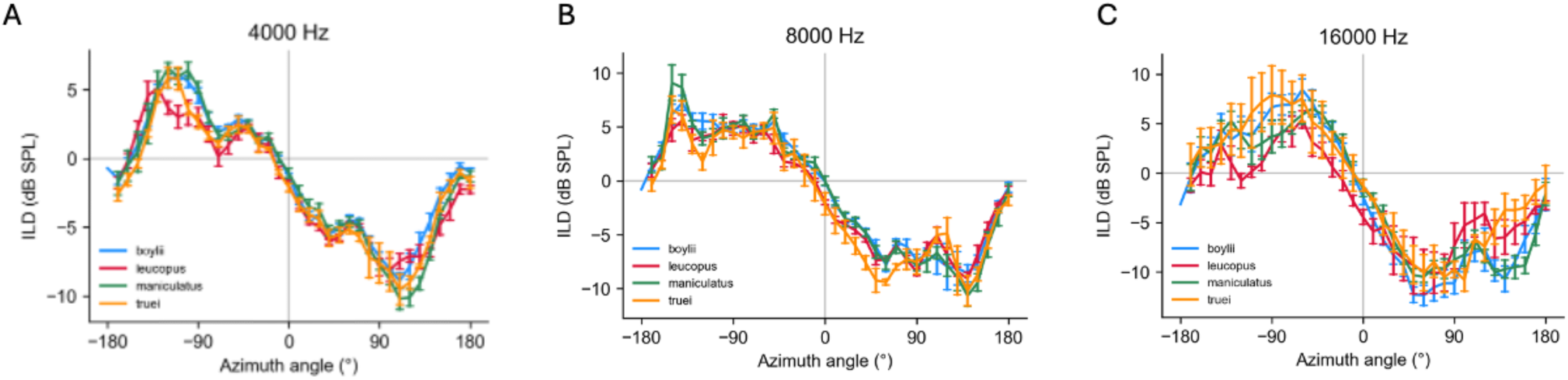
ILD per azimuth angle at 4 kHz, 8 kHz, and 16 kHz for each species. Species-wide ILD results for each azimuth angle at 4k, 8k, and 16kHz frequencies. –90° to +90° azimuth represents measurements taken in front of the animal, while +/-90° to +/-180° azimuth represents measurements taken behind the animal.

Raw traces of the HRTF data between *P. leucopus* and *P. truei* demonstrate how specific frequencies are altered based on the location of the sound source across different individuals. Specifically, the presence of variability observed around 10 kHz on the same side as the ear measured is indicative of the presence of spectral notches, which are denoted with red circles in Figure 8.

**Figure 8.**
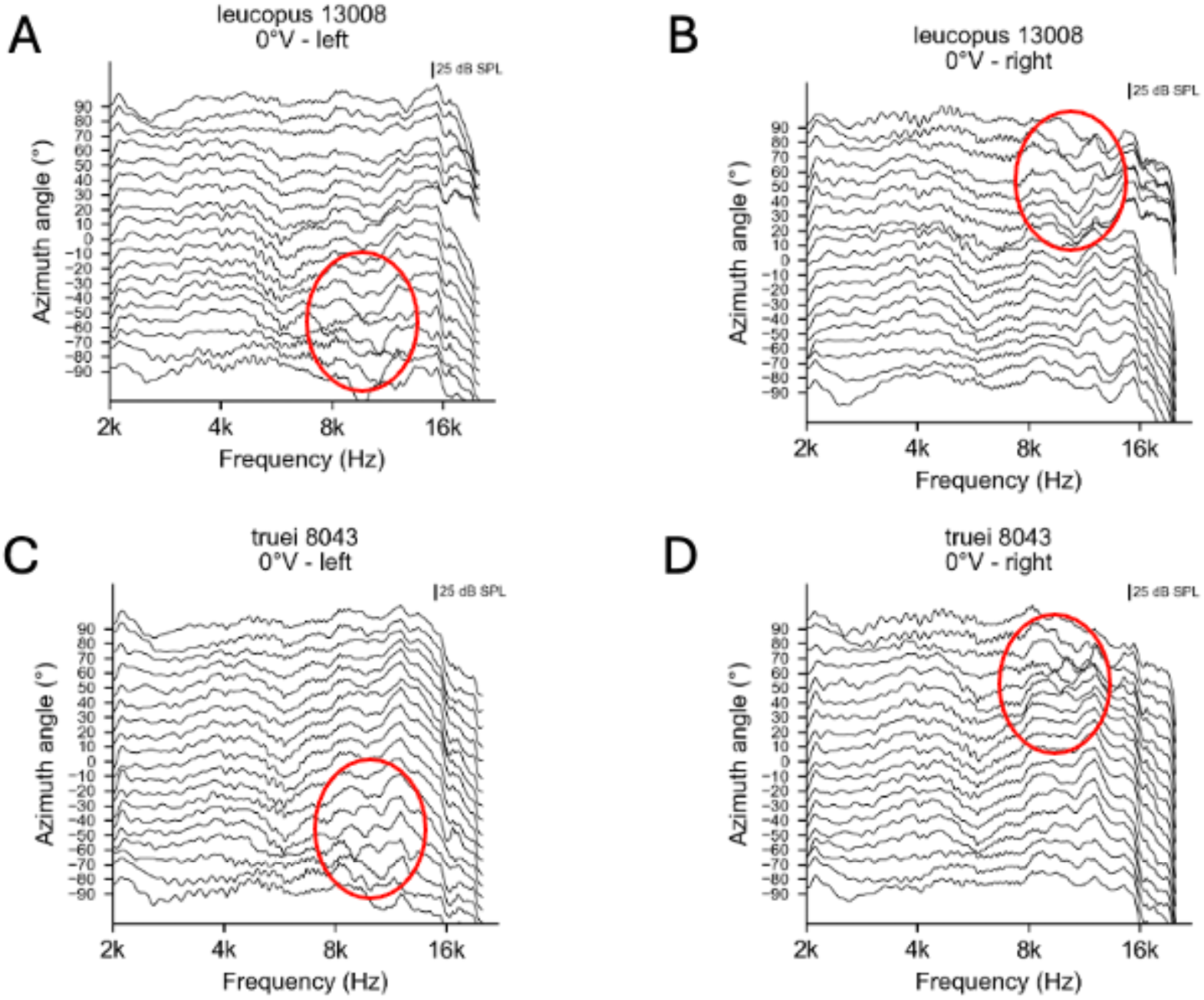
Raw HRTF traces for two individual animals. -90° in azimuth represents the left side of the animal while red circles denote variability at higher frequencies, suggesting the presence of spectral notches at ipsilateral positions in comparison to contralateral positions.

## Discussion

Previous literature describes the use of museum specimens in auditory research utilizing the skull and middle ear bones, but few researchers have performed HRTFs on zoological specimens (Rébillat et al., 2014). Works that have done so typically used taxidermy mounts, which understandably provide a more realistic biological shape and size compared to average museum specimens (Rébillat et al., 2014; Benichoux, Rébillat, and Brette, 2016). Our findings align with this rationale, as they indicate that specimen preparation and aging influence the morphology, as pinna measurements taken on preserved specimens do not reflect the measurements taken at the time of capture. Despite this, our results show significant differences in pinna length, pinna width, and IT measurements across species in ways consistent with the described difference among them. Thus, even though preservation might alter details of the relative size of soft tissue structures, these differences might disrupt within-species variation but maintain significant signal, allowing for the use, albeit limited, of these specimens in auditory research.

The results of our HRTF demonstrated consistent results across species, suggesting the differences in head and pinna size identified in our external measurements do not modify auditory cues. Even though we expect that species with the largest head and pinna (i.e., *P. boylii* or *P. truei*) would produce longer ITDs and ILDs, this has not been observed. The results of the HRTF suggest that all species investigated have their ITD and ILD cues optimized for higher frequencies, which would be consistent with an adaptation to help animals precisely locate sounds of those frequencies in their environment (Heffner and Heffner, 2008).

This correlates with what we know about the physical properties of sound, as shorter wavelengths (higher frequencies) are more affected by the head and pinna than are longer wavelengths (lower frequencies). Mice, along with many other mammals, utilize high-frequency hearing to aid in sound localization, a trait that is robustly correlated with functional head size (Heffner and Heffner, 2005). Mice rely extensively on the detection of these signals due to the mechanical restraints of their small morphology, as smaller heads do not block low-frequency signals nearly as well as higher frequencies. Thus, to have optimal binaural cues to detect the approximate location of a sound, frequencies must be high enough to be affected by the head and pinna. Though, there are exceptions to this, as some small desert mammals hear and detect especially low frequencies (below 3 kHz) to be better adapted to the sounds in their environment, while subterranean mammals have completely lost high-frequency hearing and sound localization abilities (Heffner and Heffner, 2005; Mason et al., 2017). It is possible that animals living within habitats that scatter and reflect sound more, such as within canyon slopes, have adapted by having larger pinna to optimize the available cues (Mason et al., 2017). We cannot conclude this with the current study, as HRTF results were consistent across species, but a larger sample size may provide a clearer relationship between habitat and functional implications of morphological differences.

If the results from our HRTF are correct, i.e., all species share a similar auditory profile, this could be a case of a “many-to-one mapping,” which explains functional redundancy in varying morphological structures (Wainwright et al., 2005; Thompson et al., 2017). In the case of *Peromyscus*, this could be a function of their small size, leading to functional degeneration, implying that slight changes in head and pinna sizes wouldn’t be enough to cause a functional impact. This could release pinna size to respond to other ecological pressures, such as the ones implied by Allen’s rule, which states that animals in colder environments will evolve shorter extremities, including pinna, to preserve temperature (Allen, 1877; Alhajeri et al., 2020). While this morphological trend can be observed among species (*P. truei*, which had the larger pinna are known from warmer and drier regions), we failed to see any clear geographical trend in pinna size within the most widely distributed species in our sample, *P. maniculatus* (either not shown, or supplementary material). While this could be evidence of a lack of Allen’s rule within this species, it could also be the effect of drying and aging in disturbing within-species variation. Studies with either fresh specimens or a series of preserved specimens collected and measured in a similar way might be necessary to investigate both the presence of Allen’s rule and intraspecific local adaptation in pinna size.

## Conclusions

While we provide the first evidence for the usefulness of preserved specimens in studying auditory function, our results should be interpreted with caution. While the morphology represented in preserved specimens might not represent the morphology of live animals, differences among species were observed, and they should have translated into different auditory performance. The fact that they didn’t suggest that differences of the sort we can observe in these preserved specimens do not impact the auditory cues investigated here and suggests similar performance among the investigated species. These conclusions hinge on various assumptions which should be the focus of future work. On specimen preservation, experiments should be conducted investigating not only the direct and observable effects of time on preserved specimens but also their possible role in affecting HRTF results. On auditory performance, studies including a broader range of species with different morphologies and sizes and from a wider habitat range would allow the assessment of the impact of size and environment. Furthermore, future research should explore spectral notch cues across elevations to better understand how sounds are perceived at any given point in space, rather than horizontal cues, which were studied here. By addressing these considerations, we can bridge knowledge gaps and importantly strengthen the reliability of using preserved specimens in comparative auditory research.

## Acknowledgments

We sincerely thank the Collection of Vertebrates of Oklahoma State University for loaning us their valuable research specimens and Stillwater Designs (Kicker) of Stillwater, Oklahoma, for the construction of our equipment (especially Steve Irby and Aaron Surratt). A special thanks to Peyton Williams and Cameron Miller for their help in constructing our equipment as well. Finally, thank you to Dr. Michael Reichert for the use of his sound-attenuating chamber to make HRTF measurements.

## Competing interests

No competing interests declared.

## Funding

This work was in part supported by the Oklahoma Network-Research and Mentoring for Post-Baccalaureates, NSF RaMP DBI 2216648, G.A.

## Data availability

No publicly available datasets were used throughout this work.

## Supplementary files

**Table S1.** Pairwise comparisons for pinna length across species.

| Species | Adjusted p-value |
| --- | --- |
| californicus-boylii | 0.0235 |
| gossypinus-boylii | 0.0289 |
| leucopus-boylii | 0.0000 |
| maniculatus-boylii | 0.0000 |
| truei-boylii | 0.0000 |
| gossypinus-californicus | 0.0001 |
| leucopus-californicus | 0.0000 |
| maniculatus-californicus | 0.0000 |
| truei-californicus | 0.5591 |
| leucopus-gossypinus | 0.0116 |
| maniculatus-gossypinus | 0.0000 |
| truei-gossypinus | 0.0000 |
| maniculatus-leucopus | 0.0000 |
| truei-leucopus | 0.0000 |
| truei-maniculatus | 0.0000 |

**Table S2.**
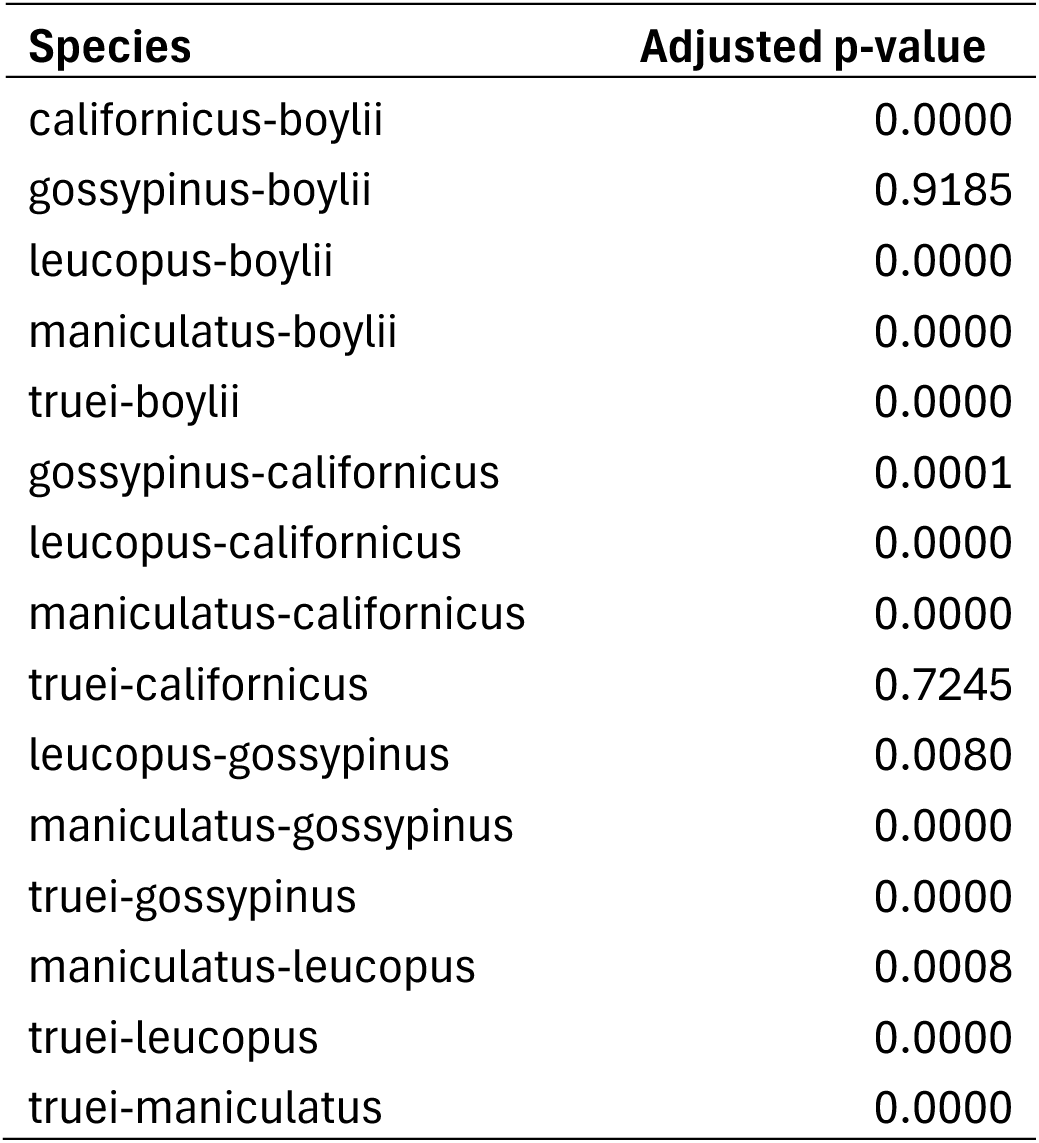
Pairwise comparisons for pinna width across species.

**Table S3.** Pairwise comparisons for inter-pinna distance across species.

| Species | Adjusted p-value |
| --- | --- |
| californicus-boylii | 0.6408 |
| gossypinus-boylii | 0.7997 |
| leucopus-boylii | 0.0000 |
| maniculatus-boylii | 0.0000 |
| truei-boylii | 0.9999 |
| gossypinus-californicus | 0.3056 |
| leucopus-californicus | 0.0000 |
| maniculatus-californicus | 0.0047 |
| truei-californicus | 0.6296 |
| leucopus-gossypinus | 0.0000 |
| maniculatus-gossypinus | 0.3008 |
| truei-gossypinus | 0.8816 |
| maniculatus-leucopus | 0.0000 |
| truei-leucopus | 0.0000 |
| truei-maniculatus | 0.0000 |

**Table S4.** Pairwise comparisons for nose-to-pinna distance across species.

| **Species** | **Adjusted p-value** |
| --- | --- |
| californicus-boylii | 0.9999 |
| gossypinus-boylii | 0.9999 |
| leucopus-boylii | 0.9995 |
| maniculatus-boylii | 0.9999 |
| truei-boylii | 0.9999 |
| gossypinus-californicus | 1.0000 |
| leucopus-californicus | 1.0000 |
| maniculatus-californicus | 0.9999 |
| truei-californicus | 1.0000 |
| leucopus-gossypinus | 1.0000 |
| maniculatus-gossypinus | 0.9999 |
| truei-gossypinus | 1.0000 |
| maniculatus-leucopus | 0.9402 |
| truei-leucopus | 1.0000 |
| truei-maniculatus | 0.9995 |

**Table S5.** Minimum and maximum mean ILD values for each species tested at 2, 4, 8, and 16 kHz. *P. boylii* n = 5, P*. leucopus* n = 5, *P. maniculatus* = 5, *P. truei* n = 3.

| 2kHz |  |  | 4kHz |  |  |
| --- | --- | --- | --- | --- | --- |
| Species | ILD min (dB) | ILD max (dB) | Species | ILD min (dB) | ILD max (dB) |
| boylli | -6.4196 | 2.5798 | boylli | -9.0702 | 6.377 |
| leucopus | -8.7834 | 2.3158 | leucopus | -8.9936 | 5.722 |
| maniculatus | -7.3558 | 1.7156 | maniculatus | -10.6142 | 7.6416 |
| truei | -6.775 | 1.473666667 | truei | -9.844666667 | 5.94 |

**Table S5. Minimum and maximum mean ILD values for each species tested at 2, 4, 8, and 16 kHz.** *P. boyllii* n = 5, *P. leucopus* n = 5, *P. maniculatus* = 5, *P. truei* n = 3
| 8kHz |  |  | 16kHz |  |  |
| --- | --- | --- | --- | --- | --- |
| Species | ILD min (dB) | ILD max (dB) | Species | ILD min (dB) | ILD max (dB) |
| boylli | -10.6512 | 8.6238 | boylli | -13.9154 | 8.7516 |
| leucopus | -9.8846 | 6.682 | leucopus | -14.2418 | 6.8718 |
| maniculatus | -11.9798 | 9.937 | maniculatus | -11.6494 | 7.4036 |
| truei | -10.67366667 | 7.991333333 | truei | -11.92 | 8.584333333 |

**Table S6.** Minimum and maximum mean ITD values for each species tested. . *P. boylii* n = 5, P*. leucopus* n = 5, *P. maniculatus* = 5, *P. truei* n = 3

| Species | ITD min (μs) | ITD max (μs) |
| --- | --- | --- |
| boylli | -335.601 | 353.741 |
| leucopus | -308.39 | 380.952 |
| maniculatus | -326.531 | 376.417 |
| truei | -332.577 | 385.488 |

## Notes

### Competing Interest Statement

The authors have declared no competing interest.

